# Alterations to cellular metabolism are linked to multiple natural parasite infections across populations of a freshwater fish

**DOI:** 10.1101/2025.05.08.652957

**Authors:** Vincent Mélançon, Sandra A. Binning, Ariane Côté, Sophie Breton

## Abstract

Exposure to environmental stressors can induce physiological responses in organisms, which can lead to population-level differences in physiological traits. However, the mechanisms underlying these responses both within individuals and among populations are often unknown. Parasite infections, in particular, are important biotic stressors that can induce a range of effects in hosts, including altered whole-organism metabolism. However, few studies have explored sub-cellular alterations to metabolic performance across both an individual infection gradient and population-level differences in infection prevalence. We compared mitochondrial enzyme activities across five distinct populations of pumpkinseed sunfish (*Lepomis gibbosus*) differing in the prevalence of cestode and trematode parasites. Overall, we found that enzymes from the OXPHOS pathway (cytochrome c oxidase, NADH dehydrogenase and coenzyme Q : cytochrome c oxidoreductase), the anaerobic pathway (lactate dehydrogenase), the TCA cycle (citrate synthase) and lipid metabolism (carnitine palmitoyl transferase) were positively correlated with parasite density with trends depending on the organ, parasite species and host population studied. Our results suggest complex and nuanced relationships among infections and host physiological performance and provide strong evidence of parasite-induced alterations to mitochondrial metabolism which may explain previously reported whole-organism responses to infections.

## Introduction

Wild animals face a wide range of environmental stressors which can trigger physiological responses that allow individual adjustment and acclimatization to these stressors [1]. These individual responses can vary depending on external factors, such as the duration of exposure, or internal factors like life stage, sex or tolerance levels [2, 3]. Indeed, substantial individual variation in physiological traits — including metabolism, thermal tolerance, and immune responsiveness — has been documented across various species and tend to be repeatable [4–7]. For example, the thermal plasticity of metabolic rates in cunners (*Tautogolabrus adspersus*) varies between individuals and is notably more repeatable in warmer waters than in colder ones [7]. Populations can also exhibit different physiological responses to a stressor, complicating the extrapolation of findings from individuals within a single population to the species as a whole. Bugg et al. [8] studied the thermal tolerance and metabolic rates of two populations of lake sturgeon (*Acipenser fulvescens*) and reported population-specific responses, with a lower thermal tolerance in the northernmost population, highlighting the role of thermal history in shaping physiological traits. Population-level differences in physiological or behavioural responses can even occur between geographically connected and closely situated populations [9–12]. This variability poses a significant challenge for conservation managers trying to predict a species’ ability to physiologically adjust to environmental perturbations. Conservation management plans typically assume that perturbations will impact metapopulations, i.e. spatially distinct populations of the same species with individual movement among populations, in similar ways [13–15]. This assumption likely stems from a lack of studies addressing population-level differences in physiological responses to stressors.

An individual’s ability to produce energy, i.e. its metabolism, is often one of the first physiological traits to adjust in response to environmental stressors. Metabolic adjustments can be made rapidly and are plastic, resulting in context-dependent variation in energy production [2, 16]. Environmental fluctuations and/or infections, for instance, have been shown to modulate physiological processes [17–19]. These modulations can be influenced by the life history of individuals. For example, a population of cavefish (*Astyanax mexicanus*) exhibits distinct immune investment patterns compared to surface-dwelling counterparts, likely due to reduced parasite diversity in the cave environment, which in turn affects their responses to bacterial infections [20]. While the mechanisms underlying population-level differences remain unclear, recent research has investigated potential subcellular physiological drivers, particularly in response to temperature variations [21, 22]. Mitochondria play a key role in organismal responses to environmental stressors, not only because of their function in energy production through oxygen consumption and cell signaling [23–26], but also due to their lesser-known crucial involvement in immune responses [27]. Mitochondria generate key immune signaling molecules, such as reactive oxygen species (ROS), and contribute to inflammation during parasite infections [28]. To produce these molecules, mitochondria adjust the activity of their metabolic pathways, which can result in altered mitochondrial metabolism [29–31]. Like organismal traits, mitochondrial dynamics exhibit plasticity at both intra- and interpopulation levels [32, 33]. For example, multiple populations of perciforms acclimated to cold temperatures tend to increase mitochondrial abundance and modulate enzyme activities, whereas populations in warmer environments decrease mitochondrial abundance and limit aerobic activities [33–35]. The plasticity of mitochondrial responses and their link to whole-organism physiology underscore the importance of including subcellular mechanisms in study designs, a practice that remains underutilized in studies of population-level responses to environmental stressors.

The effects of abiotic stressors on organismal physiology, from the subcellular to the population level, have been increasingly studied in recent years [2, 33]. In contrast, responses to biotic interactions such as predation, competition or parasitism, remain comparatively underexplored. Parasite infections, in particular, can elicit pronounced physiological responses in hosts, yet are often overlooked in studies of organismal performance traits [36]. Infections can negatively affect host health and impact survival and fitness [37–39]. Notably, parasite infections can have contradicting effects on host metabolism [37]. For example, in pumpkinseed sunfish, increasing trematode infection has been linked to reduced maximal oxygen uptake in one population, whereas a different study on another population found the opposite effect in response to infection by the same parasite species [17, 37, 40]. Such inconsistencies highlight a key limitation in current research: most studies focus on whole-organism responses within a single population and neglect subcellular mechanisms, hindering a deeper understanding of host-parasite interactions at the cellular level [37]. Discrepancies may be caused by population-specific responses driven by differences in unexamined underlying mechanisms. Existing studies linking parasite infection to mitochondrial function are limited by their narrow scope, often restricted to a single population, a single organ and/or a single parasite species [9, 17, 41, 42]. However, wild animals are typically exposed to a diversity of parasites affecting different organ systems, making broader and more integrative approaches essential for a realistic understanding of host physiological responses [43].

Co-infection, where a host is simultaneously infected by multiple parasite species, can drive differences in population-level responses to infection through priority effects and immune activation induced by the initial parasite [44–46]. In addition, parasite abundance, defined as the number of individuals of a given parasite species infecting a host, can significantly affect how hosts respond and adapt to infections [20, 47, 48]. Populations that differ in parasite richness and abundance may exhibit distinct physiological strategies. On the one hand, hosts may modulate their immune investment based on the energetic costs of immune responses relative to the burden of infection [20, 48]. Populations exposed to more parasites are hypothesized to be better “prepared”, having co-evolved with a more diverse parasite community [20, 48]. As stated in Lindstrom [48], “any immune investment will be wasted in an environment without parasites”. Populations that are used to parasite infections may favor less energetically costly adaptative immune reactions, i.e. the acquired immunity that respond in a specific manner [20, 49]. On the other hand, high parasite abundance may also reflect more productive, resource-rich environments [47, 48]. Increased resource availability can enhance host metabolic capacity, enabling greater energy allocation to immune defenses [47]. This, in turn, can select for more virulent parasites and elevated parasite abundance, prompting hosts to invest even more heavily in defense mechanisms [47]. Such feedback loops may result in higher resistance to particular parasites, but also render infection and co-infection harsher on hosts [47]. Consequently, in resource-rich environments, the fitness costs of parasitism may be amplified [47]. Considering the complex interplay between exposure, co-infection and host responses, studying mitochondrial responses to parasitic infections across environments could offer valuable insights into the subcellular mechanisms underlying whole-organism metabolic alterations, especially given mitochondria’s roles in both immune function and plastic responses to environmental stressors.

Using a freshwater fish and its helminth parasites as a model system, we aim to address the following questions: Does mitochondrial metabolism in specific host organs vary along a natural gradient of parasitic infection, and is this response consistent across different populations? We hypothesize that, due to the energetic costs associated with immune responses and the depletion of host resources caused by parasites, mitochondrial activity will be linked to parasitic infections. Building on previous findings showing that parasite infections can negatively affect whole-organism oxygen uptake in this study system [17, 18], we predict a shift toward increased anaerobic metabolism and reduced aerobic metabolism in infected individuals, with possible differences depending on the type of parasite infection [9, 17, 41]. We also expect that this relationship will differ across environments characterized by varying parasitic abundances [9].

## Methods

### Study system

Pumpkinseed sunfish (*Lepomis gibbosus*) are a small freshwater fish species native to northeastern America. *Lepomis gibbosus* is infected by multiple parasite species primarily bass tapeworms (*Proteocephalus ambloplitis*) and black spot-causing trematodes (*Uvulifer spp. & Apophalus spp.*) [50]. Both parasites have complex life cycles with *L. gibbosus* as the second intermediate host. Bass tapeworms primarily infect the liver, spleen, and gonads causing drastic tissular damage [9, 41] while black spot-causing parasites are found on the scales, gills, fins, and in muscles leading to impaired movements and immune responses which leads to the emergence of the black spots [51, 52]. *Lepomis gibbosus* can also be exposed to other parasites, such as nematodes and yellow grub trematodes (*Clinostomum marginatum*), though these occur at lower abundances and intensities [50]. The studied lake system from which *L. gibbosus* populations were sampled consists of multiple lakes with varying infection conditions [53]. More precisely, some populations experience little to no parasite infections, while others are exposed to high parasite intensities.

### Collection and procedures

To explore the links among mitochondrial metabolism and parasite infection across different populations, we studied pumpkinseed sunfish (*Lepomis gibbosus*) from five lakes (Corriveau, Croche, Cromwell, Long and Triton) around the Station de Biologie des Laurentides (SBL) in Quebec, Canada (lat. 45.98861, long. −74.00585). Despite their proximity and metapopulation structure, these lakes differ naturally in the prevalence and intensity of both blackspot and cestode infection [10, 53]. Fish were caught between July 20th and August 4th, 2021, using baited minnow traps set for 30-60 minutes. Fish were selected based on size (between 90-120 mm total length) and the presence of visible black spots such that individuals across a representative gradient of infection within a lake were chosen. Fish that did not respect these criteria were immediately released. In total, 103 fish were collected (mean total length 100.25 mm [min – max ± SE: 77.69 – 125.55 ± 0.96 mm]; mean mass 17.84g [7.06 – 35.62 ± 0.59g]). Following capture, fish were euthanized with an overdose of a 10% eugenol solution (4mL of 10% clove-oil solution / L) and immediately packed in ice to maintain tissue integrity. They were then transported to SBL’s laboratory facilities (within 2h of capture) and stored at −20°C. Fish bodies were transfer on ice to the Complexe des sciences laboratory (Université de Montréal) for further analyses.

### Sample preparation and fish dissection

Fish were dissected as in Mélançon et al. [41]. Briefly, fish bodies were partly thawed, measured using a 0-150 mm digital caliper, and weighed using an electronic balance (MSE225S, Sartorius). Organs were dissected out in the following order: heart, spleen, brain, and gills. Gills were extracted last because of the need to properly separate gills lamellae from cartilage and conjunctive tissue. Dissections were performed using a dissecting microscope (Stemi DV4, Zeiss) and on ice to maintain tissue integrity and limit light heating. Dissected organs were screened for parasites before being weighed. They were then stored in homogenization potassium phosphate buffer (100mM KP, 20mM EDTA, pH 8.0) at −80°C in cryogenic tubes at a ratio of 20 to 1 (20 times the organ wet weight). Organs were later homogenized six times for 5 sec with a homogenizer (PT1200 E, Kinematica) and placed on ice for 30 seconds between homogenizations [41]. The homogenates were used for enzymatic analyses.

Fish remains and carcasses were carefully screened for parasites (e.g. *Uvulifer spp.; Apophallus spp.; Proteocephalus ambloplitis; Clinostomum marginatum;* nematodes). Once all internal parasites were removed and counted, fish carcasses were weighed again to obtain a corrected fish mass excluding the mass of parasites [38]. Trematodes forming black cysts were only counted on one side of the fish to avoid double counting parasites on unpaired fins [54]. Fish were then disposed of according to facilities requirements.

### Enzymatic assays

Enzyme activities were measured at room temperature using a microplate reader (Mithras lab LB940, MBI). All assays were run in duplicate. Total protein content was determined using the BSA protein content assay (Sigma-aldrich kit) according to the manufacturer’s protocol. Metabolic enzyme activities [citrate synthase (CS EC 2.3.3.1) as a marker of mitochondrial metabolism, mitochondrial complexes I + III (ETS EC 7.1.1.2 and EC 7.1.1.8) and cytochrome c oxidase (CCO - Complex IV EC 7.1.1.9) as markers of aerobic metabolism, lactate dehydrogenase (LDH EC 1.1.1.27) as a marker of anaerobic metabolism, and carnitine palmitoyltransferase (CPT EC 2.3.1.21) as a marker of lipid metabolism] were assessed as follows:

CS activity was measured at 412 nm following the conversion of 5,5’dithiobis-2-nitrobenzoic acid (DTNB) into 2-nitro-5-thiobenzoic acid (TNB; ε = 14.15 mM-1 cm-1) over 6 min. The reaction solution consisted of 100 mM imidazole-HCl buffer with 0.1 mM DTNB, 0.1 mM acetyl-CoA, and 0.15 mM oxaloacetate, pH 8.0. A background without oxaloacetate was first performed followed by the final assay [41, 55].

ETS activity (combined activities of mitochondrial complexes I and III) was assessed at 490 nm over 6 min by measuring the reduction of p-iodonitrotetrazolium violet (ε = 15.9 mM-1 cm-1) using 100 mM potassium phosphate buffer with 2 mM p-iodonitrotetrazolium violet and 0.85 mM NADH, pH 8.5. Background and specificity of the reaction were calculated by running a control without tissues and a background without NADH [41, 55].

CCO activity was measured at 550 nm over 5 min by monitoring cytochrome c oxidation (ε = 18.5 mM-1 cm-1) in a 100 mM potassium phosphate buffer containing 0.05 mM oxidized cytochrome c from equine heart, 4.5 mM sodium dithionite, and 0.03 % Triton X-100, pH 8.0. Control reactions were performed by adding 10 mM sodium azide to the sample, and background activity was assessed by omitting sodium dithionite and subsequently subtracted from the measured assay activity. The absorbance ratio at 550∶565 nm was determined and if the ratio exceeded 9, the solution was used. If the ratio was below 9, the reaction solution was bubbled for 5 min, and the absorbance ratio was reassessed to reach a value of 9 [41, 55, 56].

LDH activity was measured at 340 nm over 6 min by monitoring the oxidation of NADH (ε = 6.22 mM-1 cm-1) in a 100 mM potassium phosphate buffer with 0.15 mM NADH and 0.4 mM pyruvate, pH 7.0. Background was obtained by omitting the sample [9, 56].

CPT activity was assessed at 412 nm over 5 min by measuring the conversion of DTNB) into TNB (ε = 14.15 mM-1 cm-1) in a 75 mM Tris-HCl buffer containing 1.5 mM EDTA with 0.25 mM DTNB, 0.035 mM palmitoyl-CoA, and 2 mM L-carnitine, pH 8.0. Control reactions were performed by omitting the sample [41, 56].

All enzymatic activities are expressed in U per mg of protein where U stands for 1 µmol of substrate transformed into product per minute.

### Statistical analysis

To study the link between mitochondrial enzyme activities and parasite infection, we first separately compared parasite densities (CD and BSD) among lakes. We then performed model selection for each enzymatic assay in each organ to identify the most representative model. All statistical analysis were performed using R statistical software version 4.3.2 [57].

#### Parasite infection

To compare parasite infection among individuals, individual parasite intensity (number of parasites on a host fish) divided by the total fish wet-weight (g) were used to measure three parasite densities (parasite/g) per individual: black spot density (BSD), cestode density (CD) and combined black spot and cestode density (BSCD). Densities were used to standardize parasite counts across fish size. The correlation between cestode and trematode density was assessed using Kendall’s rank correlation test as data did not follow a normal distribution. Since the correlation was low (τ= 0.04), each parasite density was included as a separate covariable in the analyses (Figure 1C). CD and BSD among lakes was compared using a non-parametric Kruskal-Wallis test followed by a post-hoc Dunn test with a Bonferroni correction.

**Figure 1:**
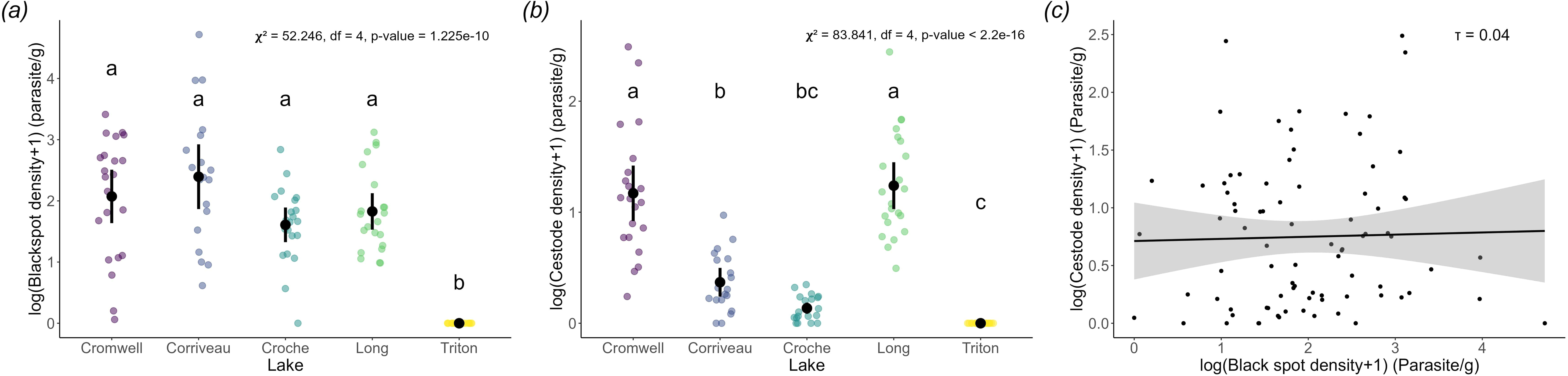
Parasite infections among different populations and their correlation. Representation of the log transformed parasite densities (parasite/g) of each lake and the correlation between parasite species. Coloured circles represent individual data, black circles represent population means and vertical lines 95% confidence intervals. Letters represent significant differences. (*a*) Infection density of black spot causing trematodes (*Uvulifer spp.* & *Apophalus spp.*). (*b*) Infection density of cestodes (*Proteocephalus ambloplitis*). (*c*) Correlation between trematode density and cestode density.

#### Enzymatic activities and parasite infection

To examine the relationship between enzymatic activities and parasite infection, we built generalized linear models using a gamma distribution with a log link which were compared using a second-order Akaike information criterion (AICc) in order to select the model that explained more of the variation without over-fitting using the MuMIn package [58]. In total, five general models were built to explain enzymatic activities differences using lake of origin and parasites densities (BSD, CD and BSCD). Models were built from most complex to least complex but always included Lake of origin and at least one infection measure (Table S1): 1) BSD, CD and lake with all interactions; 2) BSD, CD and lake without interactions between parasites densities; 3) CD and lake with all interactions; 4) BSD and lake with all interactions; 5) BSCD and lake with all interactions.

After initial data exploration, parasite densities were ln transformed to respect linear model assumptions and a value of 1 was added to parasite densities because of the presence of zeros. All explanatory variables were scaled for model analysis by subtracting the mean to all values (center) and by dividing values by their standard deviation (scale). All generalized linear models were compared using the maximum likelihood random effect estimation (ML) to allow comparison by AICc. This model selection was performed for all organs independently. When more than one model fitted the data similarly (ΔAICc of less than 2), all models were used for analysis but only the model with the lowest AICc is reported in the following section.

Model significance was assessed by comparing the models to a null model using a χ² test for variance. A similar test without the null model was used to look at the significance of each explanatory variable. Finally, confidence intervals of each estimate were used in combination with the estimate’s p-value (α = 0.05) following a post-hoc t-test to determine significant differences among lakes and slopes’ significance. 95% confidence intervals were obtained using the emmeans package [59] and were considered significant if the interval did not include zero. Post-hoc t-tests were performed with the same package. Post-hoc p-values were adjusted using the Bonferroni method.

## Results

### Parasite infection

All fish were infected except those from Lake Triton. Infected fish either had both cestode and trematode infections, or only one in rare cases (n = 7 fish). Individuals had between 0 and 846 black spots (*Uvulifer spp. & Apophalus spp*) with a median black spot density of 4.28 parasite/g (0 – 110.94 ± 1.34 parasite/g). For cestodes (*P. ambloplitis*), the median density was 0.40 parasite/g (0 – 11.06 ± 0.20 parasite/g) with infection intensity ranging from 0 to 210 cestodes. When Lake Triton fish were removed, median parasite densities were 5.29 and 0.93 parasite/g for trematodes and cestodes, respectively. Significant differences in trematode and cestode density were found among lakes (trematodes: (Figure 1A; ANOVA, χ²_4_= 52.246, p= 1.225 × 10^−10^); cestodes: Figure 1B; ANOVA, χ²_4_= 83.841, p< 2.2 × 10^−16^). For trematodes, fish from Lake Triton had a significantly different infection density than other lakes whereas for cestodes, Lake Triton had the lowest infection density, Lake Croche was similar to both Lake Triton and Corriveau and Lake Cromwell and Long had the highest cestode densities. For other parasites, different life stages of yellow grubs (*C. marginatum*; [min-max, median] 0 – 18, 0 yellow grubs) and a species of nematode (0 – 1, 0 nematodes) were detected in relatively low prevalences (respectively, 34.0% and 4.9%) and thus not included in subsequent analyses. Preliminary observations indicated that these parasites had no discernible impact on our results.

### Enzymatic activities and parasite infection

After model selection, the most parsimonious model for enzymatic activity measurements included black spot density (BSD) and the lake of origin for almost all organs and enzymes studied (Table S1). Exceptions to this were CPT activity in the heart, gills and spleen, CCO activity in the heart and spleen, ETS activity in the spleen and LDH activity in the brain and heart, where the most informative model included cestode density (CD) instead of BSD. CCO activity in the gills followed a similar model but with combined parasite densities (BSCD) as the most informative model. Some organs had multiple models within 2 AICc units and thus equally informative, but only the model with the lowest AICc will be shown in the following graphs (Tables S2-3). All models significantly explained data variations when compared to a null model (Table S2). Post-hoc results and confidence interval are reported in table S4.

Many models showed significant correlations between parasite density and enzymatic activity. In many instances, a positive correlation between BSD and enzymatic activity was found. Indeed, this correlation was found for ETS in the heart (ANOVA, χ²_1_= 0.772, *p*= 0.043; figure 2C) and spleen (ANOVA, χ²_1_= 5.538, *p*= 0.00021), for CCO in the gills (ANOVA, χ²_1_= 1.165, *p*= 1.476 × 10^−08^), for CS in the heart (ANOVA, χ²_1_= 7.536, *p*= 1.102 × 10^−12^; figure 4C), for CPT in the spleen (ANOVA, χ²_1_= 1.156, *p*= 0.020), and for LDH in the gills (ANOVA, χ²_1_= 6.375, *p*= 2.056 × 10^−15^; figure 6B), heart (ANOVA, χ²_1_= 10.199, *p*< 2.2 × 10^−16^), and spleen (ANOVA, χ²_1_= 15.873, *p*< 2.2 × 10^−16^; figure 6D). Similar positive correlations between CD and enzymatic activity were also found for CCO in the spleen (ANOVA, χ²_1_= 18.431, *p*< 2.2 × 10^−16^; figure 3D) and for CPT in the heart (ANOVA, χ²_1_= 0.562, *p*= 0.044; figure 5C) and between BSCD and enzymatic activity for ETS in the spleen (ANOVA, χ²_1_= 5.899, *p*= 0.00013), for CCO in the gills (ANOVA, χ²_1_= 0.914, *p*= 5.405 × 10^−07^; figure 3B) and for LDH in the spleen (ANOVA, χ²_1_= 16.344, *p*< 2.2 × 10^−16^). On the other hand, negative correlations between BSD and enzymatic activity were discovered for CCO in the brain (ANOVA, χ²_1_= 0.741, *p*= 0.0017; figure 3A) and CS in the spleen (ANOVA, χ²_1_= 1.234, *p*= 0.0079; figure 4D) and between CD and enzymatic activity for CCO in the gills (ANOVA, χ²_1_= 0.188, *p*= 0.024), for LDH in the brain (ANOVA, χ²_1_= 0.705, *p*= 0.0057; figure 6A) and in the heart (ANOVA, χ²_1_= 0.368, *p*= 0.047; figure 6C).

**Figure 2:**
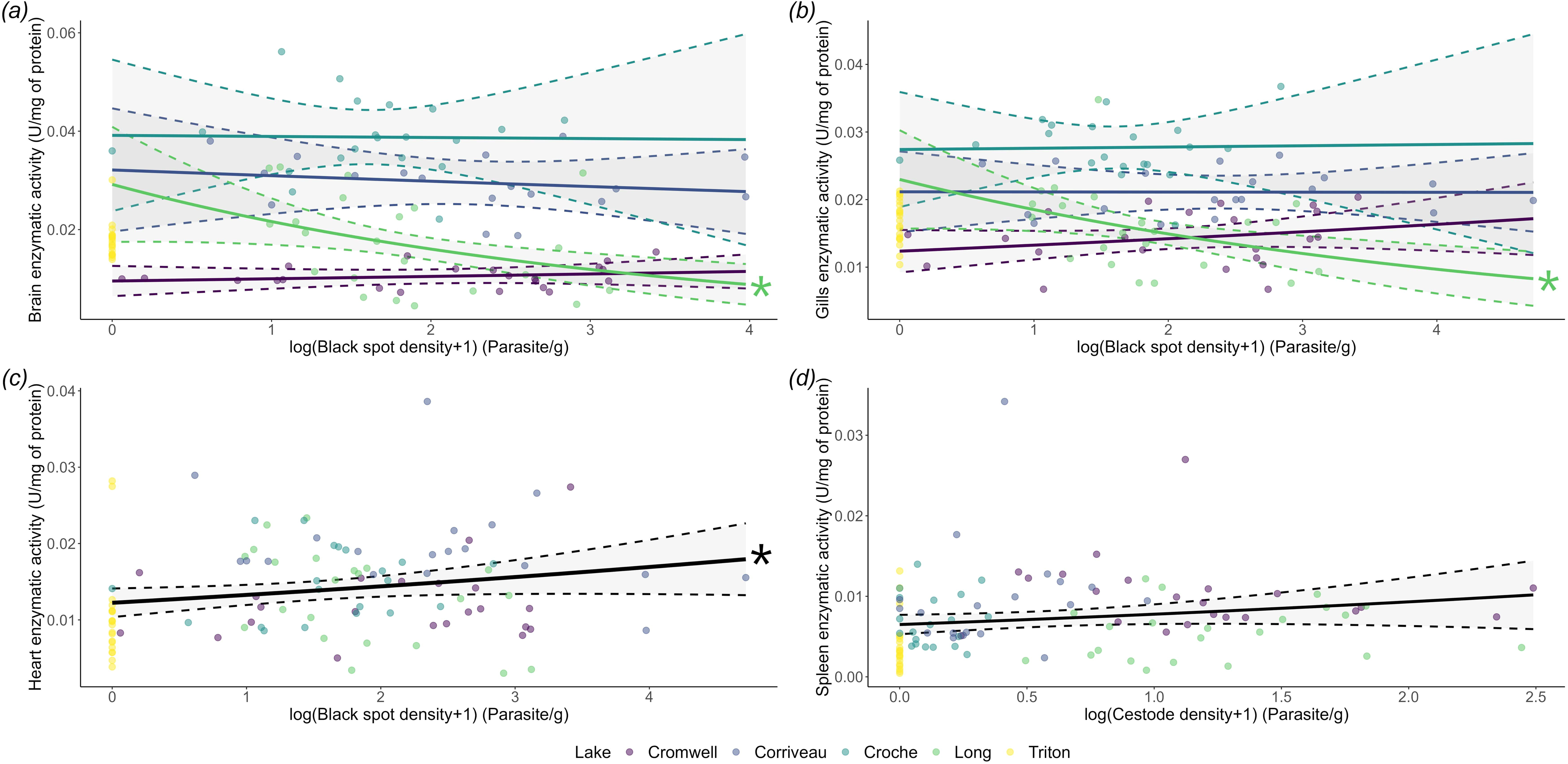
Enzymatic activity of ETS for all organs in relation to parasite density. Link between log transformed parasite density (parasite/g) and enzymatic activities (U • mg of protein^-1^). Coloured circles represent enzymatic activities of all individuals, solid lines the linear regression and dotted lines 95% confidence intervals. Asterix represent significant slopes. (*a*) Brain enzymatic activity in relation to BSD. (*b*) Gills enzymatic activity in relation to BSD. (*c*) Heart enzymatic activity in relation to BSD. (*d*) Spleen enzymatic activity in relation to CD.

**Figure 3:**
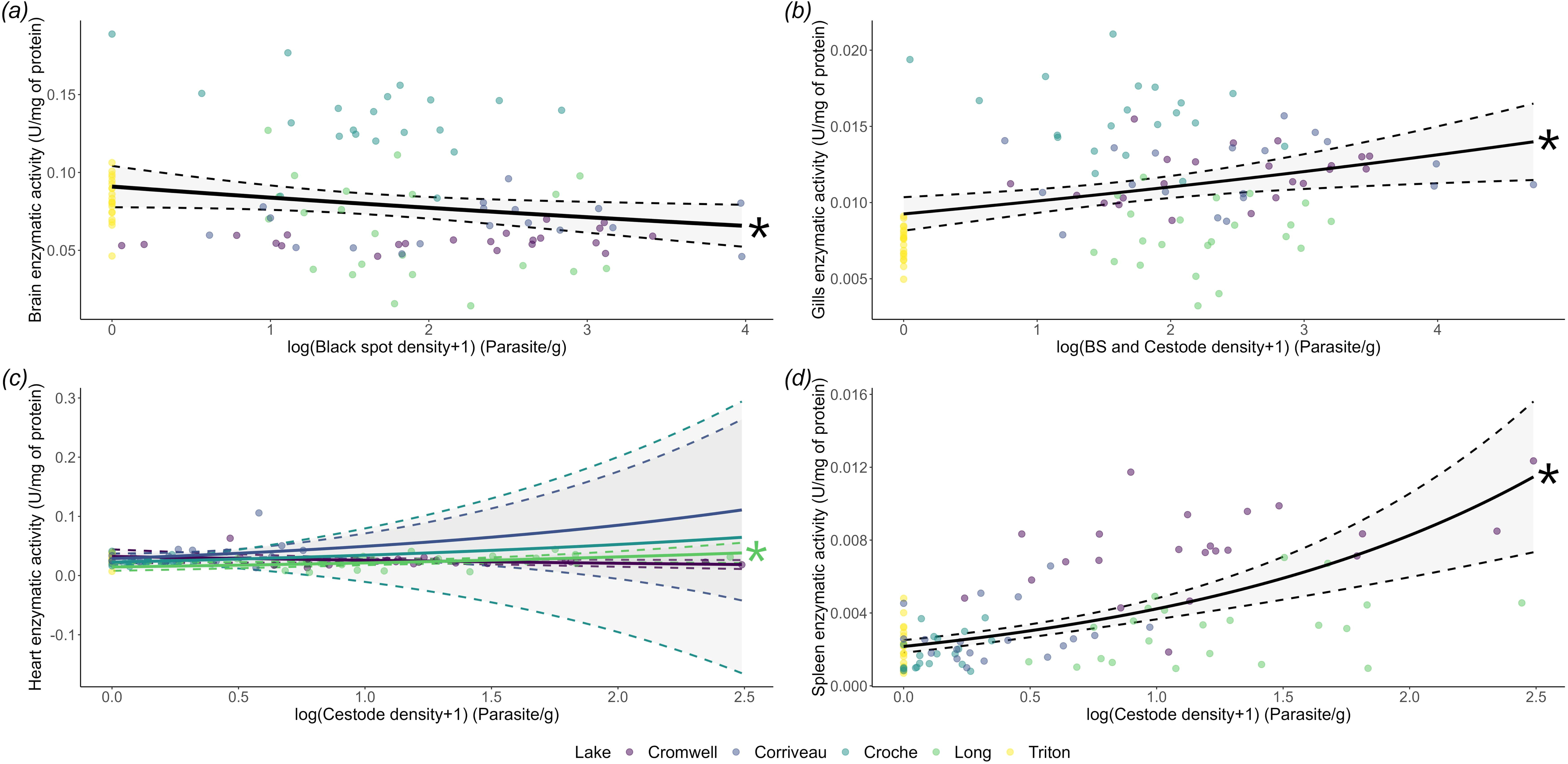
Enzymatic activity of CCO for all organs in relation to parasite density. Link between log transformed parasite density (parasite/g) and enzymatic activities (U • mg of protein^-1^). Coloured circles represent enzymatic activities of all individuals, solid lines the linear regression and dotted lines 95% confidence intervals. Asterix represent significant slopes. (*a*) Brain enzymatic activity in relation to BSD. (*b*) Gills enzymatic activity in relation to BSCD. (*c*) Heart enzymatic activity in relation to CD. (*d*) Spleen enzymatic activity in relation to CD.

**Figure 4:**
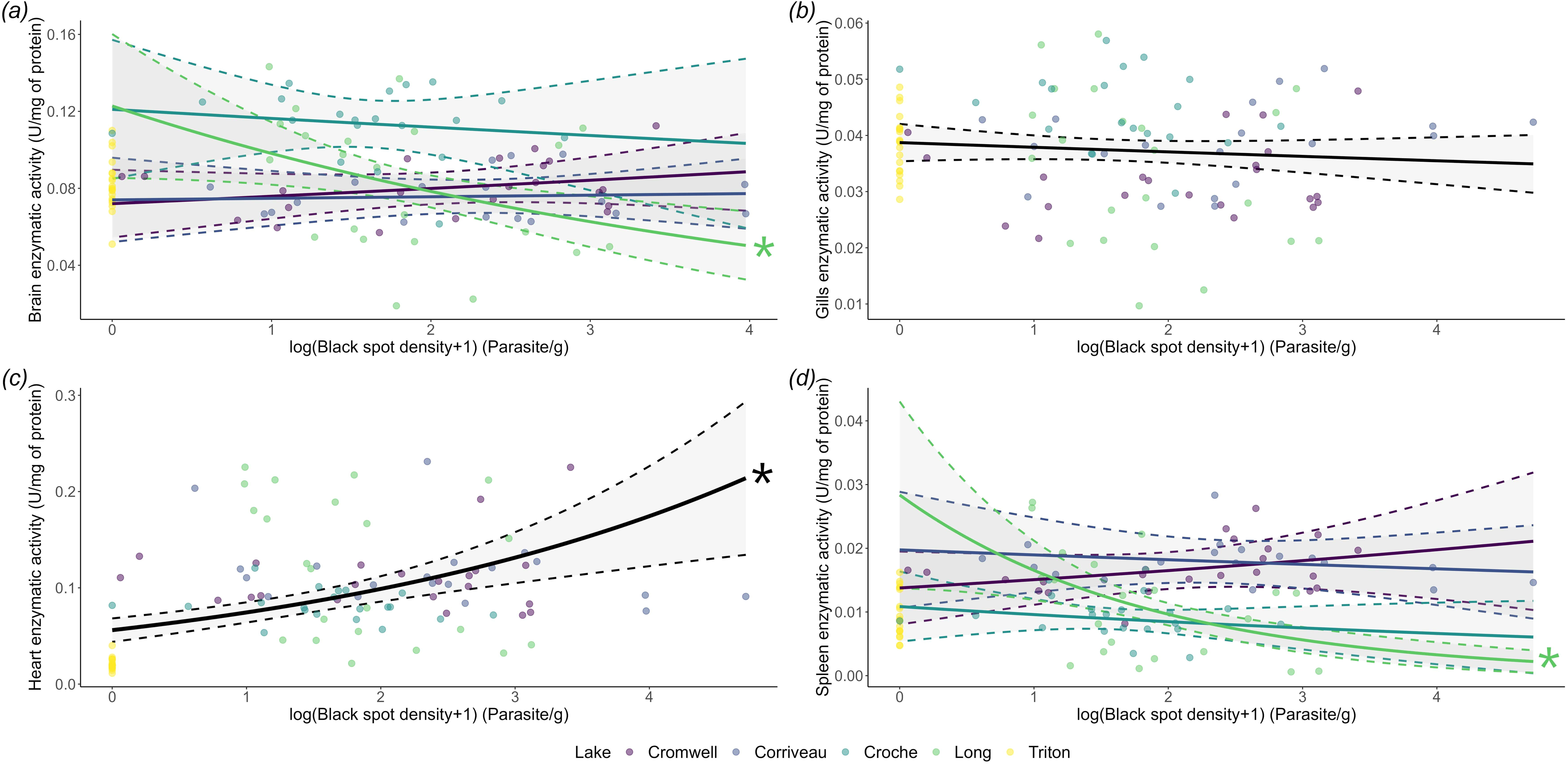
Enzymatic activity of CS for all organs in relation to parasite density. Link between log transformed parasite density (parasite/g) and enzymatic activities (U • mg of protein^-1^). Coloured circles represent enzymatic activities of all individuals, solid lines the linear regression and dotted lines 95% confidence intervals. Asterix represent significant slopes. (*a*) Brain enzymatic activity in relation to BSD. (*b*) Gills enzymatic activity in relation to BSD. (*c*) Heart enzymatic activity in relation to BSD. (*d*) Spleen enzymatic activity in relation to BSD.

**Figure 5:**
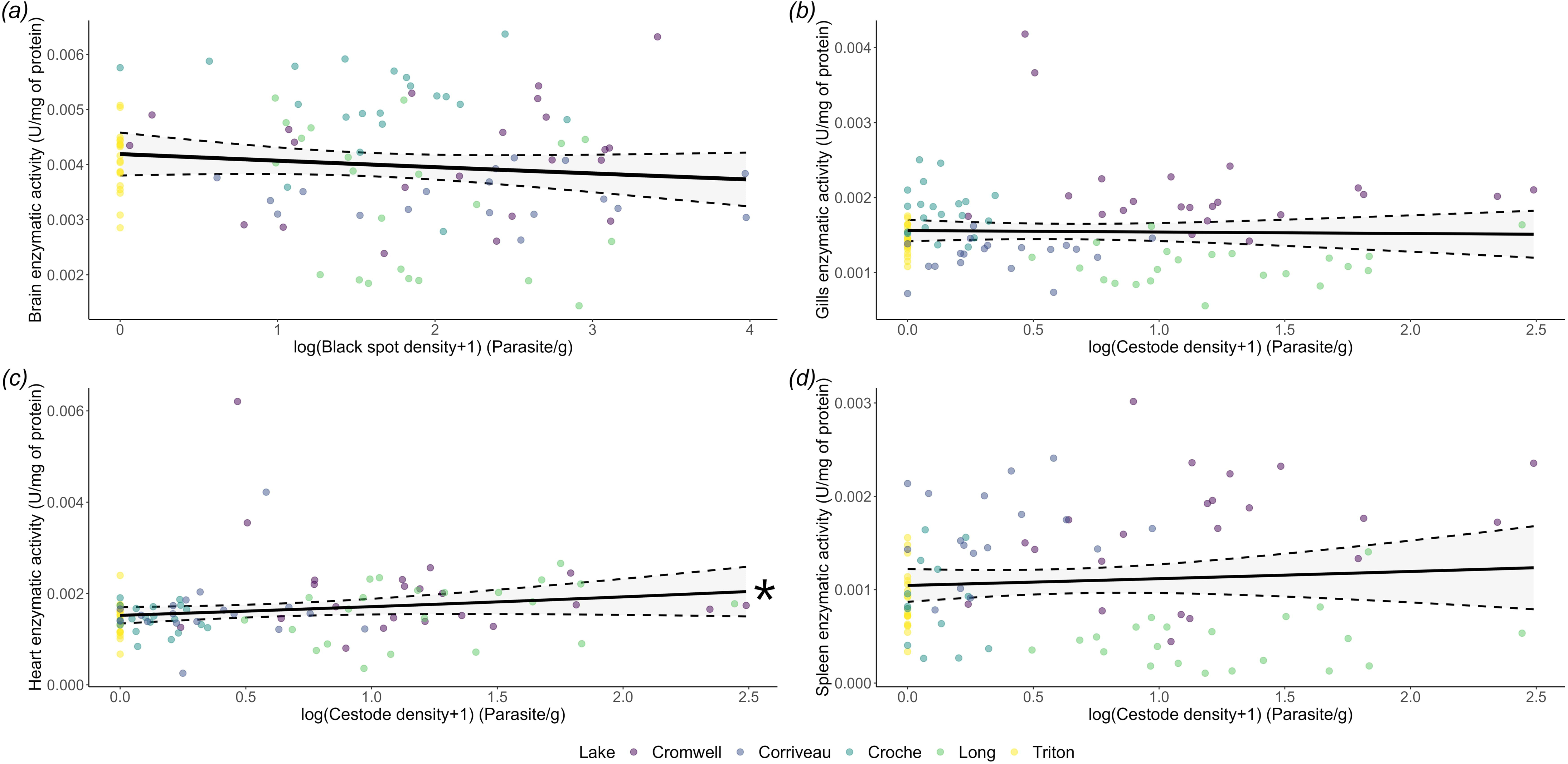
Enzymatic activity of CPT for all organs in relation to parasite density. Link between log transformed parasite density (parasite/g) and enzymatic activities (U • mg of protein^-1^). Coloured circles represent enzymatic activities of all individuals, solid lines the linear regression and dotted lines 95% confidence intervals. Asterix represent significant slopes. (*a*) Brain enzymatic activity in relation to BSD. (*b*) Gills enzymatic activity in relation to CD. (*c*) Heart enzymatic activity in relation to CD. (*d*) Spleen enzymatic activity in relation to CD.

**Figure 6:**
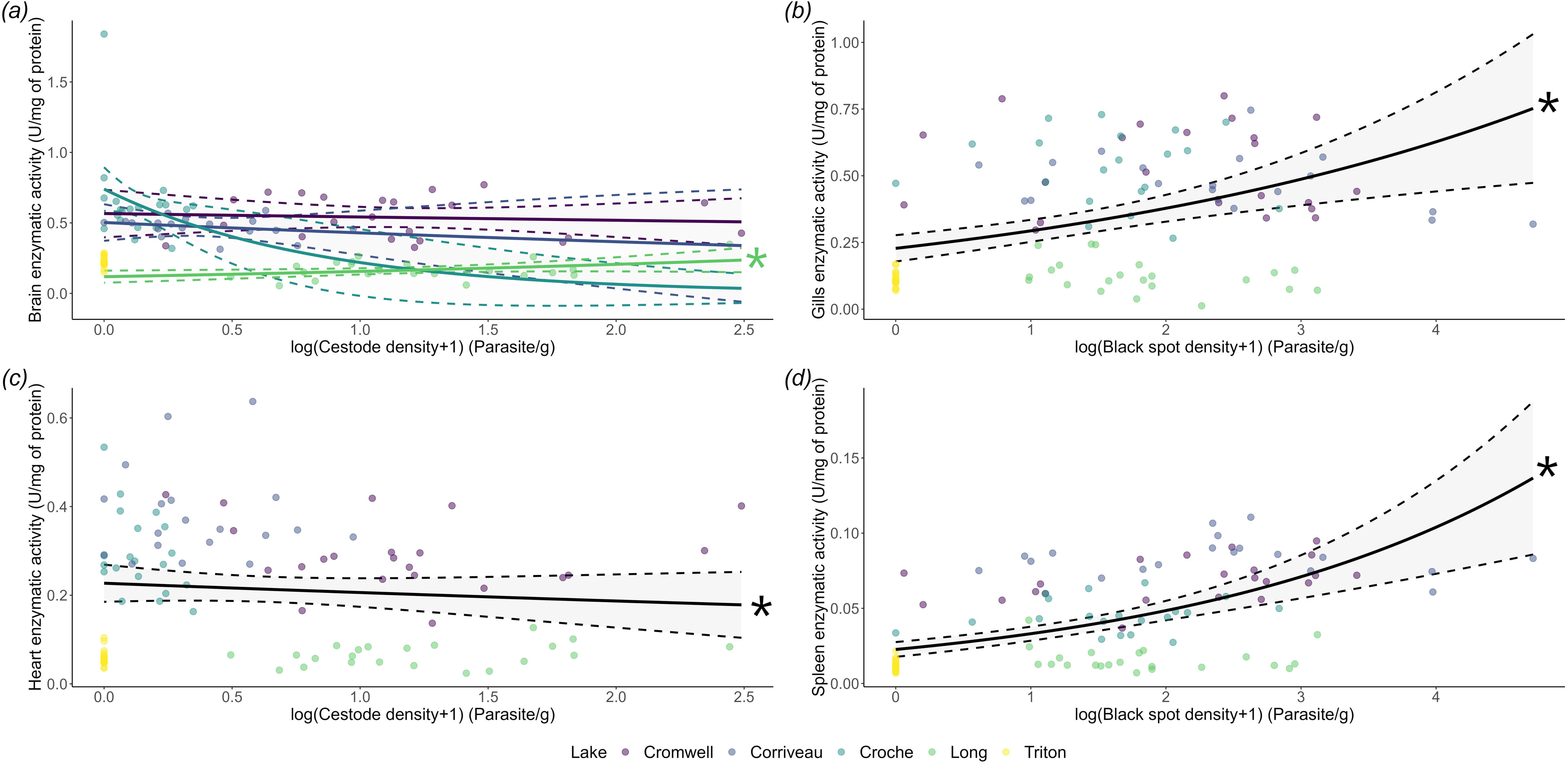
Enzymatic activity of LDH for all organs in relation to parasite density. Link between log transformed parasite density (parasite/g) and enzymatic activities (U • mg of protein^-1^). Coloured circles represent enzymatic activities of all individuals, solid lines the linear regression and dotted lines 95% confidence intervals. Asterix represent significant slopes. (*a*) Brain enzymatic activity in relation to CD. (*b*) Gills enzymatic activity in relation to BSD. (*c*) Heart enzymatic activity in relation to CD. (*d*) Spleen enzymatic activity in relation to BSD.

All enzyme activities showed an effect of the lake of origin (Table S3), with the exception of CPT in the heart (ANOVA, χ²_4_= 1.295, *p*= 0.052). A summary of the comparisons can be found in Table S4. In general, fish from Lake Triton and Lake Long tended to exhibit lower enzyme activities, while fish from Lake Cromwell almost always showed higher activities of all enzymes. For ETS, fish from Lake Corriveau had higher activities. For CCO, fish from Lake Croche had higher activities in the brain and gills, while fish from Lake Cromwell showed higher activity in the spleen. Fish from Lake Triton and Long, on the other hand, showed lower activity in the gills. Regarding CS, fish from Lake Long and Cromwell showed higher activities in both the heart and gills, while fish from Lake Croche and Triton had lower activities in these organs. For CPT, fish from Lake Cromwell and Croche showed higher activities in the brain, gills and spleen, while fish from Lake Long showed lower activities. Finally, for LDH, fish from Lake Long and Triton showed lower activity in the heart, gills and spleen while fish from Lake Cromwell and Corriveau had higher activities in the same three organs.

In some cases, the interaction between parasite density and lake of origin was found to be significant. For ETS, the interaction between BSD and lake was significant in the brain (ANOVA, χ²_3_= 0.867, *p*= 0.041; figure 2A) and gills (ANOVA, χ²_3_= 0.548, *p*= 0.039; figure 2B). For CCO, the interaction between CD and lake was significant in the heart (ANOVA, χ²_3_= 1.37, *p*= 0.021; figure 3C). For CS, the interaction between BSD and lake was significant in the brain (ANOVA, χ²_3_= 0.554, *p*= 0.027; figure 4A) and spleen (ANOVA, χ²_3_= 2.674, *p*= 0.0016; figure 4D). For CPT, the interaction was never significant. For LDH, the interaction between CD and lake was significant in the brain (ANOVA, χ²_3_= 0.795, *p*= 0.035; figure 6A). When an interaction was significant, the relationship between Lake Long and parasite density was significantly different from that of Lake Cromwell, except for CS in the spleen, where it was also significantly different from Lake Corriveau, and for LDH in the brain, where Lake Long showed no differences compared to the other Lakes (Table S4). Moreover, the slope for Lake Long was significant in all instances when using confidence intervals, while the slopes for the other lakes were never significant (Table S4).

## Discussion

We examined the activity of mitochondrial enzymes in pumpkinseed sunfish naturally infected by parasites across five populations with varying levels of infection prevalence. As expected, we found significant correlations between enzymatic activities and parasite densities in most of the organs studied, with variations observed depending on the type of parasite studied. Moreover, we identified population-specific differences in the relationship between enzymatic activities and parasite infections. We expected to observe different effects on hosts depending on the parasite species, but no such differences were observed in this study, contrary to previous findings [41]. Our results show that parasite infections are correlated to sub-cellular physiological traits confirming the role of mitochondria in host responses to infection and suggesting a link with whole-organism metabolic responses to parasite infections. They also point out the importance of studying multiple populations as it can highlight population responses that can differ from species-wide trends.

### Correlations between parasite densities and enzyme activities

We hypothesized a relationship between enzymatic activities and host parasite density [9, 17, 41]. Specifically, we expected a decrease in aerobic enzyme activities and an increase in anaerobic activities when parasite abundances were higher. Our findings revealed significant correlations between parasite densities and enzyme activities, with variations observed depending on the organ studied. Contrary to what was expected, complex IV (CCO) showed increased activity in the gills and spleen, and a decrease in the brain while complexes I and III (ETS) showed increased activity in the heart and spleen. These findings suggest an overall increase of the OXPHOS pathway, the central ATP production of the aerobic metabolism, when fish are more heavily infected [23]. Mitochondria are crucial in immune responses, playing roles in inflammation signalling, immune cell activation, and defence mechanisms [27]. ETS and CCO play a key role in maintaining the proton gradient across mitochondrial membranes, which is essential for ATP synthesis [23, 24, 60]. The observed increase in ETS and CCO activity in more parasitized fish suggests an enhanced proton gradient, leading to greater ATP production [23, 24, 60]. Higher ATP production may be needed to support responses to parasite infections such as immune cell activation, damage repair or inflammation [27]. While ATP production is the main function of mitochondria, the observed increase in OXPHOS activities, especially in the spleen, could be mainly linked to increased ATP demand but also with increased immune signalling through reactive oxygen species (ROS), in response to parasite infection [27]. Our results somewhat contradict findings exploring bacterial or viral infections, where hosts tend to decrease activity of OXPHOS complexes after the inflammation reaction [29]. This difference could be explained by the fact that we studied ongoing infections and not acute reactions. Longterm infections could be linked with reduced inflammation, the mechanism thought to diminish OXPHOS activities once triggered [29], therefore leading to increased OXPHOS activity. The increased costs could be prominently linked to tissue repair and adaptative immune responses, which are linked to higher OXPHOS activities [27]. This could also point out fundamental differences between bacterial/viral and metazoan infections. This could potentially show that latter infections are more costly and could lead to bigger physiological responses.

The observed increase in OXPHOS activity must be supported by upstream pathways. Indeed, we also found a positive correlation between parasite infection and CS (TCA cycle) activity in both the heart and spleen. Additionally, a slight positive correlation between lipid metabolism (CPT) and parasite infection was observed in the same organs. Both pathways (i.e. TCA cycle and lipid metabolism) assist OXPHOS to produce ATP [61–63]. Increases in these enzyme activities are therefore most likely a response to the increase in OXPHOS activity. Additionally, many intermediates of the TCA cycle play key roles in immune responses, acting as signalling molecules with important regulatory effects on inflammation and immune cell activation [27]. Lipid metabolism supports the TCA cycle by providing essential molecules [61, 62]. An increase of the TCA cycle either because of an increased OXPHOS activity or immune signalling could lead to increased lipid metabolism. It is also possible that parasite infections induce a state of starvation, resulting in increased lipid metabolism activity. Indeed, parasites can deplete host resources and affect nutrient absorption [64], potentially leading to starvation or malnutrition. On another note, CS is thought to give indication of mitochondrial abundance [65]. If an increase in CS is linked to an increase in mitochondrial abundance, it could explain why OXPHOS and CPT activities also increased. Mitochondrial abundances can change following acclimation to certain environmental stressors and such a response to parasite infection could be plausible [33, 34]. Indeed, mitochondrial morphology has been shown to react to intracellular parasite infections [66, 67].

Anaerobic metabolism (LDH activity) also showed significant correlations with parasite infections. Specifically, positive correlations were observed in the gills, heart, and spleen in response to parasite infection, while the brain exhibited a negative correlation. Although aerobic metabolism is the primary pathway for ATP production due to its higher yield [68], anaerobic metabolism, specifically anaerobic glycolysis mediated by LDH, can supplement ATP production when necessary [69]. This pathway produces ATP alternatively to OXPHOS using by-products of the TCA cycle [69]. The increased LDH activity in the heart and spleen could meet the increased ATP demand of immune responses, while also ensuring proper OXPHOS functions, a key role of the anaerobic pathway [69].

The observed decrease in LDH activity in the brain confirms previous results [42]. Nadler et al. [42] found that brain anaerobic potential decreased following parasite infection and hypothesized this decrease in LDH activity to be a mechanism behind behaviour-manipulating parasites. While we cannot confirm if such behaviour manipulation is present in our study system, black spot-causing trematodes have been linked to alterations in host behaviour following an experimental infection consistent with results suggesting decreased responsiveness to aerial attacks [18, 70]. We also hypothesize that this response could protect hosts as an accumulation of lactate in the brain would be detrimental and could cause damage. Additionally, the decreased CCO activity in the brain could have a similar protective role. Parasites tend to rely on aerobic mechanisms when oxygen is available [71], therefore depleting the host’s resources. By reducing oxygen demand in certain organs, an individual could prevent oxygen deficiency while maintaining normal functions and avoiding ROS accumulation. Organs of the circulatory system (i.e. gills and heart) might be better equipped to resist the adverse effects of ROS, while the spleen requires these molecules for immune responses. The circulatory organs may be more solicitated to facilitate circulation of immune cells to infection sites, thereby preventing ROS or damaged cells from accumulating in other areas, which could explain the upregulation of enzyme activities in these organs.

Overall, our findings suggest a mitochondrial chain of reactions that enable an organism to respond to infection by producing more energy potentially enabling them to activate immune cells, produce signalling molecules (ROS, TCA intermediates), repair cell and tissue damage caused either by the infection or by immune activity, or maintain activity levels despite increased energetic needs. The increase in OXPHOS activity seems to be supported by increased activities of the TCA cycle and lipid metabolism and is accompanied by increased activity of the anaerobic metabolism.

### Parasite species

Parasite species can have different effects on their host depending on their general ecology (i.e. life cycle, targeted tissues, transmission, activity) [41]. We found that black spot-causing trematodes were more often correlated with enzymatic activities than cestodes and better accounted for the variation in the data. Surprisingly, both types of parasites exhibited positive and negative correlations with enzyme activities, although positive correlations were more common. Also, in some rare instances, combined parasite infections were linked with enzyme activities and explained the observed variation.

It was previously suggested that trematodes might have less detrimental direct effects on their hosts with associated metabolic responses linked primarily to immune reactions during the acute phase of infection [41]. However, our findings show that both types of infection induce similar trends, i.e. increased mitochondrial enzyme activities. Although trematode density in this study spanned a wide range, with many individuals showing either high or low infection levels, cestode infection densities were generally lower, and only three individuals had greater than 7 parasite/g cestodes. This narrower infection range for cestodes may have limited our ability to detect similar trends as seen with trematodes. Hence, we cannot conclude whether the effects of the two parasite species differ or are similar, although our data suggest the latter. This limitation could be overcome by controlling the infection rates of both parasites through experimental infections as a future research direction.

### Differences among populations

Despite their geographic proximity, the different populations studied experience different levels of exposure to parasites. As reported, trematodes causing black spot infections showed similar levels of infection in most lakes, except in Lake Triton, which exhibited no infection at all. On the contrary, cestode infection varied in intensity among lakes, with Lake Cromwell and Lake Long exhibiting the highest levels, while Lake Croche and Lake Corriveau showed similarly low infection intensities. These exposure differences could lead to different physiological responses [20, 48, 72]. Our research also identified several enzyme activities that exhibited variations in their relationships between the lake of origin and parasite infections. In general, fish from Lake Long responded differently than those from Lake Cromwell and significantly to parasite infections. Finally, in almost all cases, significant absolute differences in enzyme activity, independent of infection status, were observed across lakes.

Population-level differences in the lakes we studied could be due to life-history differences in immune investment versus reproduction and growth, driven by differing levels of parasite exposure [20, 47, 48, 73]. Evidence suggests that fish from these populations exhibit differences in growth rates, with fish from Lake Long and Lake Triton growing and maturing faster than fish from other sampled lakes (Beauchamp, unpublished data). Thus, the reduced enzyme activities observed in fish from Lake Long may reflect energetic trade-offs unique to this population. Additionally, general LDH activity levels in fish from Lake Long were lower than those of other populations, which could explain why their responses to infection differ. Fish from Lake Long may have been exposed to parasites for a longer time, as the population of the final hosts (smallmouth bass – *Micropterus dolomieu*) has reportedly been established in this lake for a longer period. In contrast, the introduction of *M. dolomieu* into Lake Cromwell occurred only recently (within the last 20 years) following a natural dispersal event. Fish from Lake Long, accustomed to parasite exposure, could be less impacted by their presence [20] and show smaller physiological stress, while the more naïve population from Lake Cromwell may mount a stronger immune response to combat infections [74]. These life-history differences could explain why trends between Lake Long and Lake Cromwell differ significantly. Moreover, absolute differences in enzymatic activities (i.e. independent of parasite densities) show that population have different base levels in some cases, highlighting the possibility of population-specific physiological adaptations.

## Conclusion

In light of these results, we report a correlation between parasite infections and mitochondrial enzyme activities. Contrary to our initial predictions, we observed an increase in both aerobic and anaerobic energy production pathways. While some differences among populations were identified, our results predominantly suggest that the physiological responses appear to be similar species wide. Comparing populations remains challenging, and numerous factors could explain our results. Nevertheless, we emphasize the importance of incorporating multiple populations in study designs, as some differences may emerge. Overall, trends encompassing all populations should still be considered and interpreted, as they reflect the shared exposure of *L. gibbosus* to the same parasites and may help in conservation management efforts. However, inter-population differences can drive certain patterns in opposite directions, giving a more comprehensive understanding of host-parasite interactions. These differences do not hinder our interpretations but rather highlight the complexity of host-parasite dynamics, which should be examined across multiple biological scales.

## Aknowledgments

We acknowledge the traditional lands of the Kanien’kehà:ka, Omàmiwinini, and Anishinabewaki First Nations on which the field and laboratory work for this project took place. We thank Gabriel Lanthier for logistic support at the SBL and his team. We thank Amélie Papillon, Alexandra Kack, Maryane Gradito and Marie Levet for their help during fish sampling and useful conversations. We also thank Ariane Pouliot-Drouin, Sofia Sabbagh, Thomas McNeil, Tommy Pepin and Tristan Briand for their assistance during lab experiments.

## Ethics

All experiments were approved by the Université de Montréal’s animal care committee and the ministère de l’Environnement, de la Lutte contre les Changements Climatiques et de la Faune et des Parcs (Comité de Déontologie de l’Expérimentation sur les Animaux; Permit number: 21-028, Collection permit number: 2021-05-18-1833-15-S-P).

## Use of Artificial Intelligence (AI) and AI-assisted technologies

AI technologies were not use in the preparation of this article

## Data, code and materials

Database and code can be found on figshare through these links:

https://doi.org/10.6084/m9.figshare.28532672

https://doi.org/10.6084/m9.figshare.28532633

