## Supplemental Table 1 for "Alterations to cellular metabolism are linked to multiple natural parasite infections across populations of a freshwater fish"

| Model number | Response variable | Explanatory variable(s) | Interactions | Random factors | Model type |
| --- | --- | --- | --- | --- | --- |
| 1 | EA | CD  BSD  Lake | CD:BSD  CD:Lake  CD :BSD :Lake |  | Generalized linear model |
| 2 | EA | CD  BSD  Lake | CD:Lake  BSD:Lake |  | Generalized linear model |
| 3 | EA | CD  Lake | CD:Lake |  | Generalized linear model |
| 4 | EA | BSD  Lake | BSD:Lake |  | Generalized linear model |
| 5 | EA | BSCD  Lake | BSCD:Lake |  | Generalized linear model |
