## Supplemental Table 2 for "Alterations to cellular metabolism are linked to multiple natural parasite infections across populations of a freshwater fish"

| Enzyme | Organ | Model | Deviance | DF | Pr(>Chi) |
| --- | --- | --- | --- | --- | --- |
| ETS | Brain | BSD.log + Lake + BSD.log:Lake | 22.16 | 8 | **< 2.2e-16** |
|  | Heart | BSD.log + Lake + BSD.log:Lake | 6.1743 | 8 | **7.189e-05** |
|  | Gills | BSD.log + Lake + BSD.log:Lake | 6.6837 | 8 | **< 2.2e-16** |
|  | Spleen | BSD.log + Lake + BSD.log:Lake | 12.332 | 8 | **0.0001628** |
|  |  | CD.log + Lake + CD.log:Lake | 12.439 | 8 | **0.0001809** |
|  |  | BS_Cest_D.log + Lake + BS_Cest_D.log:Lake | 12.229 | 8 | **0.0001874** |
| CS | Brain | BSD.log + Lake + BSD.log:Lake | 2.7364 | 8 | **3.481e-07** |
|  | Heart | BSD.log + Lake + BSD.log:Lake | 33.864 | 8 | **< 2.2e-16** |
|  | Gills | BSD.log + Lake + BSD.log:Lake | 1.7031 | 8 | **0.0003459** |
|  | Spleen | BSD.log + Lake + BSD.log:Lake | 11.234 | 8 | **6.958e-11** |
| CPT | Brain | BSD.log + Lake + BSD.log:Lake | 2.7887 | 8 | **5.901e-08** |
|  | Heart | CD.log + Lake + CD.log:Lake | 2.7063 | 8 | **0.01193** |
|  | Gills | CD.log + Lake + CD.log:Lake | 6.4461 | 8 | **< 2.2e-16** |
|  | Spleen | BSD.log + Lake + BSD.log:Lake | 21.079 | 8 | **< 2.2e-16** |
|  |  | CD.log + Lake + CD.log:Lake | 21.335 | 8 | **< 2.2e-16** |
| CCO | Brain | BSD.log + Lake + BSD.log:Lake | 11.037 | 8 | **< 2.2e-16** |
|  | Heart | CD.log + Lake + CD.log:Lake | 3.7156 | 8 | **0.0008719** |
|  | Gills | BSD.log + Lake + BSD.log:Lake | 8.2873 | 8 | **< 2.2e-16** |
|  |  | CD.log + Lake + CD.log:Lake | 8.2914 | 8 | **< 2.2e-16** |
|  |  | BS_Cest_D.log + Lake + BS_Cest_D.log:Lake | 8.3046 | 8 | **< 2.2e-16** |
|  | Spleen | CD.log + Lake + CD.log:Lake | 32.029 | 8 | **< 2.2e-16** |
| LDH | Brain | CD.log + Lake + CD.log:Lake | 26.537 | 8 | **< 2.2e-16** |
|  | Heart | BSD.log + Lake + BSD.log:Lake | 57.024 | 8 | **< 2.2e-16** |
|  |  | CD.log + Lake + CD.log:Lake | 57.046 | 8 | **< 2.2e-16** |
|  | Gills | BSD.log + Lake + BSD.log:Lake | 46.604 | 8 | **< 2.2e-16** |
|  | Spleen | BSD.log + Lake + BSD.log:Lake | 54.789 | 8 | **< 2.2e-16** |
|  |  | BS_Cest_D.log + Lake + BS_Cest_D.log:Lake | 54.776 | 8 | **< 2.2e-16** |
