## Supplemental Table 3 for "Alterations to cellular metabolism are linked to multiple natural parasite infections across populations of a freshwater fish"

| Enzyme | Organ | Model | Covariables/Factors | DF | Deviance | Pr(>Chi) |
| --- | --- | --- | --- | --- | --- | --- |
| ETS | Brain | BSD.log + Lake + BSD.log:Lake | BSD.log | 1 | 0.0127 | 0.72858 |
|  |  |  | Lake | 4 | 21.2803 | **<2e-16** |
|  |  |  | BSD.log:Lake | 3 | 0.8672 | **0.04113** |
|  | Heart | BSD.log + Lake + BSD.log:Lake | BSD.log | 1 | 0.7724 | **0.0433518** |
|  |  |  | Lake | 4 | 4.0101 | **0.0002903** |
|  |  |  | BSD.log:Lake | 3 | 1.3919 | 0.0614039 |
|  | Gills | BSD.log + Lake + BSD.log:Lake | BSD.log | 1 | 0.0159 | 0.62248 |
|  |  |  | Lake | 4 | 6.1194 | **<2e-16** |
|  |  |  | BSD.log:Lake | 3 | 0.5484 | **0.03906** |
|  | Spleen | BSD.log + Lake + BSD.log:Lake | BSD.log | 1 | 5.5384 | **0.0002076** |
|  |  |  | Lake | 4 | 6.4130 | **0.0031087** |
|  |  |  | BSD.log:Lake | 3 | 0.3805 | 0.8144244 |
|  |  | CD.log + Lake + CD.log:Lake | CD.log | 1 | 1.1915 | 0.08802 |
|  |  |  | Lake | 4 | 10.7872 | **2.694e-05** |
|  |  |  | CD.log:Lake | 3 | 0.4606 | 0.77105 |
|  |  | BS_Cest_D.log + Lake + BS_Cest_D.log:Lake | BS_Cest_D.log | 1 | 5.8986 | **0.000132** |
|  |  |  | Lake | 4 | 6.0405 | **0.004775** |
|  |  |  | BS_Cest_D.log:Lake | 3 | 0.2897 | 0.869022 |
| CS | Brain | BSD.log + Lake + BSD.log:Lake | BSD.log | 1 | 0.04007 | 0.41629 |
|  |  |  | Lake | 4 | 2.14209 | **3.981e-07** |
|  |  |  | BSD.log:Lake | 3 | 0.55424 | **0.02748** |
|  | Heart | BSD.log + Lake + BSD.log:Lake | BSD.log | 1 | 7.5364 | **1.102e-12** |
|  |  |  | Lake | 4 | 25.3375 | **<2e-16** |
|  |  |  | BSD.log:Lake | 3 | 0.9897 | 0.08387 |
|  | Gills | BSD.log + Lake + BSD.log:Lake | BSD.log | 1 | 0.06298 | 0.3021924 |
|  |  |  | Lake | 4 | 1.21862 | **0.0003808** |
|  |  |  | BSD.log:Lake | 3 | 0.42151 | 0.0680491 |
|  | Spleen | BSD.log + Lake + BSD.log:Lake | BSD.log | 1 | 1.2335 | **0.007933** |
|  |  |  | Lake | 4 | 7.3266 | **1.778e-08** |
|  |  |  | BSD.log:Lake | 3 | 2.6742 | **0.001592** |
| CPT | Brain | BSD.log + Lake + BSD.log:Lake | BSD.log | 1 | 0.10144 | 0.1811 |
|  |  |  | Lake | 4 | 6.0888 | **1.916e-08** |
|  |  |  | BSD.log:Lake | 3 | 5.7672 | 0.1288 |
|  | Heart | CD.log + Lake + CD.log:Lake | CD.log | 1 | 0.56220 | **0.04357** |
|  |  |  | Lake | 4 | 1.29501 | 0.05223 |
|  |  |  | CD.log:Lake | 3 | 0.84909 | 0.10447 |
|  | Gills | CD.log + Lake + CD.log:Lake | CD.log | 1 | 0.0070 | 0.6884 |
|  |  |  | Lake | 4 | 6.2119 | **<2e-16** |
|  |  |  | CD.log:Lake | 3 | 0.2271 | 0.1588 |
|  | Spleen | BSD.log + Lake + BSD.log:Lake | BSD.log | 1 | 1.1561 | **0.02045** |
|  |  |  | Lake | 4 | 19.5308 | **<2e-16** |
|  |  |  | BSD.log:Lake | 3 | 0.3921 | 0.61013 |
|  |  | CD.log + Lake + CD.log:Lake | CD.log | 1 | 0.1807 | 0.3581 |
|  |  |  | Lake | 4 | 21.0882 | **<2e-16** |
|  |  |  | CD.log:Lake | 3 | 0.0658 | 0.9585 |
| CCO | Brain | BSD.log + Lake + BSD.log:Lake | BSD.log | 1 | 0.7418 | **0.001654** |
|  |  |  | Lake | 4 | 9.8316 | **<2e-16** |
|  |  |  | BSD.log:Lake | 3 | 0.4640 | 0.102637 |
|  | Heart | CD.log + Lake + CD.log:Lake | CD.log | 1 | 0.03309 | 0.627309 |
|  |  |  | Lake | 4 | 2.31243 | **0.002444** |
|  |  |  | CD.log:Lake | 3 | 1.37006 | **0.020711** |
|  | Gills | BSD.log + Lake + BSD.log:Lake | BSD.log | 1 | 1.1646 | **1.476e-08** |
|  |  |  | Lake | 4 | 7.0380 | **<2e-16** |
|  |  |  | BSD.log:Lake | 3 | 0.0847 | 0.5061 |
|  |  | CD.log + Lake + CD.log:Lake | CD.log | 1 | 0.1876 | **0.02353** |
|  |  |  | Lake | 4 | 8.0775 | **<2e-16** |
|  |  |  | CD.log:Lake | 3 | 0.0263 | 0.86874 |
|  |  | BS_Cest_D.log + Lake + BS_Cest_D.log:Lake | BS_Cest_D.log | 1 | 0.9137 | **5.405e-07** |
|  |  |  | Lake | 4 | 7.3037 | **<2e-16** |
|  |  |  | BS_Cest_D.log:Lake | 3 | 0.0873 | 0.4938 |
|  | Spleen | CD.log + Lake + CD.log:Lake | CD.log | 1 | 18.4314 | **<2e-16** |
|  |  |  | Lake | 4 | 12.8499 | **5.742e-11** |
|  |  |  | CD.log:Lake | 3 | 0.7479 | 0.3716 |
| LDH | Brain | CD.log + Lake + CD.log:Lake | CD.log | 1 | 0.7052 | **0.005696** |
|  |  |  | Lake | 4 | 25.0367 | **<2e-16** |
|  |  |  | CD.log:Lake | 3 | 0.7951 | **0.034816** |
|  | Heart | BSD.log + Lake + BSD.log:Lake | BSD.log | 1 | 10.199 | **<2e-16** |
|  |  |  | Lake | 4 | 46.307 | **<2e-16** |
|  |  |  | BSD.log:Lake | 3 | 0.518 | 0.1424 |
|  |  | CD.log + Lake + CD.log:Lake | CD.log | 1 | 0.368 | **0.04704** |
|  |  |  | Lake | 4 | 56.020 | **<2e-16** |
|  |  |  | CD.log:Lake | 3 | 0.658 | 0.07004 |
|  | Gills | BSD.log + Lake + BSD.log:Lake | BSD.log | 1 | 6.375 | **2.056e-15** |
|  |  |  | Lake | 4 | 39.838 | **<2e-16** |
|  |  |  | BSD.log:Lake | 3 | 0.391 | 0.2764 |
|  | Spleen | BSD.log + Lake + BSD.log:Lake | BSD.log | 1 | 15.873 | **<2e-16** |
|  |  |  | Lake | 4 | 38.674 | **<2e-16** |
|  |  |  | BSD.log:Lake | 3 | 0.243 | 0.4909 |
|  |  | BS_Cest_D.log + Lake + BS_Cest_D.log:Lake | BS_Cest_D.log | 1 | 16.344 | **<2e-16** |
|  |  |  | Lake | 4 | 38.205 | **<2e-16** |
|  |  |  | BS_Cest_D.log:Lake | 3 | 0.227 | 0.5177 |
