## Supplemental Table 4 for "Alterations to cellular metabolism are linked to multiple natural parasite infections across populations of a freshwater fish"

| Enzyme | Organ | Model | Analysis | Covariables/Factors | Confidence interval | p-value |
| --- | --- | --- | --- | --- | --- | --- |
| ETS | Brain | BSD.log+  Lake+  BSD.log:Lake | Trends | BSD.log:LakeLo | **-0.564 : -0.0999** |  |
|  |  |  |  | BSD.log:LakeCm | -0.107 : 0.2100 |  |
|  |  |  |  | BSD.log:LakeCo | -0.220 : 0.1375 |  |
|  |  |  |  | BSD.log:LakeCr | -0.263 : 0.2506 |  |
|  |  |  |  | BSD.log:lakeTr | NA |  |
|  |  |  | Contrasts | Lo – Cm |  | **0.0478** |
|  |  |  |  | Lo – Co |  | 0.3085 |
|  |  |  |  | Lo – Cr |  | 0.3885 |
|  |  |  |  | Lo – Tr |  | NA |
|  |  |  |  | Cm – Co |  | 1.000 |
|  |  |  |  | Cm – Cr |  | 1.000 |
|  |  |  |  | Cm – Tr |  | NA |
|  |  |  |  | Co – Cr |  | 1.000 |
|  |  |  |  | Co – Tr |  | NA |
|  |  |  |  | Cr - Tr |  | NA |
|  | Heart | BSD.log+  Lake+  BSD.log:Lake | Trends | BSD.log | -0.001568688 : 0.1818108 |  |
|  |  |  | Contrasts | Lo – Cm |  | 1.000 |
|  |  |  |  | Lo – Co |  | **0.0224** |
|  |  |  |  | Lo – Cr |  | 1.000 |
|  |  |  |  | Lo – Tr |  | 0.8977 |
|  |  |  |  | Cm – Co |  | **0.0061** |
|  |  |  |  | Cm – Cr |  | 1.000 |
|  |  |  |  | Cm – Tr |  | 1.000 |
|  |  |  |  | Co – Cr |  | 0.3094 |
|  |  |  |  | Co – Tr |  | **0.0029** |
|  |  |  |  | Cr - Tr |  | 0.1468 |
|  | Gills | BSD.log+  Lake+  BSD.log:Lake | Trends | BSD.log:LakeLo | **-0.4226 : -0.0563** |  |
|  |  |  |  | BSD.log:LakeCm | -0.0478 : 0.2021 |  |
|  |  |  |  | BSD.log:LakeCo | -0.1220 : 0.1201 |  |
|  |  |  |  | BSD.log:LakeCr | -0.1954 : 0.2101 |  |
|  |  |  |  | BSD.log:lakeTr | NA |  |
|  |  |  | Contrasts | Lo – Cm |  | **0.0337** |
|  |  |  |  | Lo – Co |  | 0.2015 |
|  |  |  |  | Lo – Cr |  | 0.4565 |
|  |  |  |  | Lo – Tr |  | NA |
|  |  |  |  | Cm – Co |  | 1.000 |
|  |  |  |  | Cm – Cr |  | 1.000 |
|  |  |  |  | Cm – Tr |  | NA |
|  |  |  |  | Co – Cr |  | 1.000 |
|  |  |  |  | Co – Tr |  | NA |
|  |  |  |  | Cr - Tr |  | NA |
|  | Spleen | BSD.log+  Lake+  BSD.log:Lake | Trends | BSD.log | **0.116641 : 0.3754771** |  |
|  |  |  | Contrasts | Lo – Cm |  | **0.0251** |
|  |  |  |  | Lo – Co |  | 0.0932 |
|  |  |  |  | Lo – Cr |  | 1.000 |
|  |  |  |  | Lo – Tr |  | 1.000 |
|  |  |  |  | Cm – Co |  | 1.000 |
|  |  |  |  | Cm – Cr |  | 0.5549 |
|  |  |  |  | Cm – Tr |  | **0.0095** |
|  |  |  |  | Co – Cr |  | 1,000 |
|  |  |  |  | Co – Tr |  | **0.0419** |
|  |  |  |  | Cr - Tr |  | 0.4048 |
|  |  | CD.log+  Lake+  CD.log:Lake | Trends | CD.log | -0.02738688 : 0.2632973 |  |
|  |  |  | Contrasts | Lo – Cm |  | **0.0181** |
|  |  |  |  | Lo – Co |  | 0.1570 |
|  |  |  |  | Lo – Cr |  | 1.000 |
|  |  |  |  | Lo – Tr |  | 1.000 |
|  |  |  |  | Cm – Co |  | 1.000 |
|  |  |  |  | Cm – Cr |  | 1.000 |
|  |  |  |  | Cm – Tr |  | **0.0333** |
|  |  |  |  | Co – Cr |  | 1.000 |
|  |  |  |  | Co – Tr |  | **0.0011** |
|  |  |  |  | Cr - Tr |  | 0.1048 |
|  |  | BS_Cest_D.log+  Lake+  BS_Cest_D.log:Lake | Trends | BS_Cest_D.log | **0.125968 : 0.3890044** |  |
|  |  |  | Contrasts | Lo – Cm |  | **0.0232** |
|  |  |  |  | Lo – Co |  | 0.0701 |
|  |  |  |  | Lo – Cr |  | 1.000 |
|  |  |  |  | Lo – Tr |  | 1.000 |
|  |  |  |  | Cm – Co |  | 1.000 |
|  |  |  |  | Cm – Cr |  | 0.7051 |
|  |  |  |  | Cm – Tr |  | **0.0393** |
|  |  |  |  | Co – Cr |  | 1.000 |
|  |  |  |  | Co – Tr |  | 0.0819 |
|  |  |  |  | Cr - Tr |  | 0.4969 |
| CS | Brain | BSD.log+  Lake+  BSD.log:Lake | Trends | BSD.log:LakeLo | **-0.4250 : -0.0726** |  |
|  |  |  |  | BSD.log:LakeCm | -0.0625 : 0.1779 |  |
|  |  |  |  | BSD.log:LakeCo | -0.1236 : 0.1476 |  |
|  |  |  |  | BSD.log:LakeCr | -0.2390 : 0.1510 |  |
|  |  |  |  | BSD.log:lakeTr | NA |  |
|  |  |  | Contrasts | Lo – Cm |  | **0.0320** |
|  |  |  |  | Lo – Co |  | 0.1319 |
|  |  |  |  | Lo – Cr |  | 0.7512 |
|  |  |  |  | Lo – Tr |  | NA |
|  |  |  |  | Cm – Co |  | 1.000 |
|  |  |  |  | Cm – Cr |  | 1.000 |
|  |  |  |  | Cm – Tr |  | NA |
|  |  |  |  | Co – Cr |  | 1.000 |
|  |  |  |  | Co – Tr |  | NA |
|  |  |  |  | Cr - Tr |  | NA |
|  | Heart | BSD.log+  Lake+  BSD.log:Lake | Trends | BSD.log | **0.1685842 : 0.4623488** |  |
|  |  |  | Contrasts | Lo – Cm |  | 1.000 |
|  |  |  |  | Lo – Co |  | 1.000 |
|  |  |  |  | Lo – Cr |  | **0.0118** |
|  |  |  |  | Lo – Tr |  | **< 0.0001** |
|  |  |  |  | Cm – Co |  | 1.000 |
|  |  |  |  | Cm – Cr |  | **0.0454** |
|  |  |  |  | Cm – Tr |  | **< 0.0001** |
|  |  |  |  | Co – Cr |  | 0.0129 |
|  |  |  |  | Co – Tr |  | **< 0.0001** |
|  |  |  |  | Cr - Tr |  | **< 0.0001** |
|  | Gills | BSD.log+  Lake+  BSD.log:Lake | Trends | BSD.log | -0.07242575 : 0.02417911 |  |
|  |  |  | Contrasts | Lo – Cm |  | 1.000 |
|  |  |  |  | Lo – Co |  | 1.000 |
|  |  |  |  | Lo – Cr |  | **0.0092** |
|  |  |  |  | Lo – Tr |  | 1.000 |
|  |  |  |  | Cm – Co |  | 0.2653 |
|  |  |  |  | Cm – Cr |  | **0.0010** |
|  |  |  |  | Cm – Tr |  | 1.000 |
|  |  |  |  | Co – Cr |  | 0.9769 |
|  |  |  |  | Co – Tr |  | 1.000 |
|  |  |  |  | Cr - Tr |  | 0.7912 |
|  | Spleen | BSD.log+  Lake+  BSD.log:Lake | Trends | BSD.log:LakeLo | **-0.897 : -0.299** |  |
|  |  |  |  | BSD.log:LakeCm | -0.104 : 0.304 |  |
|  |  |  |  | BSD.log:LakeCo | -0.243 : 0.152 |  |
|  |  |  |  | BSD.log:LakeCr | -0.468 : 0.195 |  |
|  |  |  |  | BSD.log:lakeTr | NA |  |
|  |  |  | Contrasts | Lo – Cm |  | **0.0014** |
|  |  |  |  | Lo – Co |  | **0.0173** |
|  |  |  |  | Lo – Cr |  | 0.2572 |
|  |  |  |  | Lo – Tr |  | NA |
|  |  |  |  | Cm – Co |  | 1.000 |
|  |  |  |  | Cm – Cr |  | 1.000 |
|  |  |  |  | Cm – Tr |  | NA |
|  |  |  |  | Co – Cr |  | 1.000 |
|  |  |  |  | Co – Tr |  | NA |
|  |  |  |  | Cr - Tr |  | NA |
| CPT | Brain | BSD.log+  Lake+  BSD.log:Lake | Trends | BSD.log | -0.08666591 : 0.02199022 |  |
|  |  |  | Contrasts | Lo – Cm |  | **0.0237** |
|  |  |  |  | Lo – Co |  | 1.000 |
|  |  |  |  | Lo – Cr |  | **< 0.0001** |
|  |  |  |  | Lo – Tr |  | 0.7139 |
|  |  |  |  | Cm – Co |  | 0.1709 |
|  |  |  |  | Cm – Cr |  | 0.0924 |
|  |  |  |  | Cm – Tr |  | 1.000 |
|  |  |  |  | Co – Cr |  | **0.0001** |
|  |  |  |  | Co – Tr |  | 1.000 |
|  |  |  |  | Cr - Tr |  | 0.0696 |
|  | Heart | CD.log+  Lake+  CD.log:Lake | Trends | CD.log | -0.01136419 : 0.1654451 |  |
|  |  |  | Contrasts | Lo – Cm |  | 0.4883 |
|  |  |  |  | Lo – Co |  | 1.000 |
|  |  |  |  | Lo – Cr |  | 1.000 |
|  |  |  |  | Lo – Tr |  | 1.000 |
|  |  |  |  | Cm – Co |  | 1.000 |
|  |  |  |  | Cm – Cr |  | 0.3041 |
|  |  |  |  | Cm – Tr |  | 0.3367 |
|  |  |  |  | Co – Cr |  | 1.000 |
|  |  |  |  | Co – Tr |  | 1.000 |
|  |  |  |  | Cr - Tr |  | 1.000 |
|  | Gills | CD.log+  Lake+  CD.log:Lake | Trends | CD.log | -0.07339909 : 0.05847611 |  |
|  |  |  | Contrasts | Lo – Cm |  | **< 0.0001** |
|  |  |  |  | Lo – Co |  | 1.000 |
|  |  |  |  | Lo – Cr |  | **< 0.0001** |
|  |  |  |  | Lo – Tr |  | 0.2472 |
|  |  |  |  | Cm – Co |  | **< 0.0001** |
|  |  |  |  | Cm – Cr |  | 0.6282 |
|  |  |  |  | Cm – Tr |  | **0.0004** |
|  |  |  |  | Co – Cr |  | **< 0.0001** |
|  |  |  |  | Co – Tr |  | 0.5129 |
|  |  |  |  | Cr - Tr |  | **0.0062** |
|  | Spleen | BSD.log+  Lake+  BSD.log:Lake | Trends | BSD.log | -0.006352491 : 0.222283 |  |
|  |  |  | Contrasts | Lo – Cm |  | **< 0.0001** |
|  |  |  |  | Lo – Co |  | **< 0.0001** |
|  |  |  |  | Lo – Cr |  | **0.0033** |
|  |  |  |  | Lo – Tr |  | **0.0437** |
|  |  |  |  | Cm – Co |  | 1.000 |
|  |  |  |  | Cm – Cr |  | **0.0016** |
|  |  |  |  | Cm – Tr |  | **0.0041** |
|  |  |  |  | Co – Cr |  | **0.0030** |
|  |  |  |  | Co – Tr |  | **0.0095** |
|  |  |  |  | Cr - Tr |  | 1.000 |
|  |  | CD.log+  Lake+  CD.log:Lake | Trends | CD.log | -0.0729213 : 0.1634417 |  |
|  |  |  | Contrasts | Lo – Cm |  | **< 0.0001** |
|  |  |  |  | Lo – Co |  | **< 0.0001** |
|  |  |  |  | Lo – Cr |  | **0.0006** |
|  |  |  |  | Lo – Tr |  | **0.0002** |
|  |  |  |  | Cm – Co |  | 1.000 |
|  |  |  |  | Cm – Cr |  | 0.8875 |
|  |  |  |  | Cm – Tr |  | 1.000 |
|  |  |  |  | Co – Cr |  | **0.0117** |
|  |  |  |  | Co – Tr |  | **0.0143** |
|  |  |  |  | Cr - Tr |  | 1.000 |
| CCO | Brain | BSD.log+  Lake+  BSD.log:Lake | Trends | BSD.log | **-0.1780452 : -0.002279192** |  |
|  |  |  | Contrasts | Lo – Cm |  | 1.000 |
|  |  |  |  | Lo – Co |  | 1.000 |
|  |  |  |  | Lo – Cr |  | **< 0.0001** |
|  |  |  |  | Lo – Tr |  | 0.5631 |
|  |  |  |  | Cm – Co |  | 0.4544 |
|  |  |  |  | Cm – Cr |  | **< 0.0001** |
|  |  |  |  | Cm – Tr |  | 0.0701 |
|  |  |  |  | Co – Cr |  | **< 0.0001** |
|  |  |  |  | Co – Tr |  | 1.000 |
|  |  |  |  | Cr - Tr |  | **< 0.0001** |
|  | Heart | CD.log+  Lake+  CD.log:Lake | Trends | CD.log:LakeLo | **0.0307 : 0.4654** |  |
|  |  |  |  | CD.log:LakeCm | -0.3243 : 0.0426 |  |
|  |  |  |  | CD.log:LakeCo | -0.0710 : 0.7622 |  |
|  |  |  |  | CD.log:LakeCr | -0.7071 : 1.2439 |  |
|  |  |  |  | CD.log:lakeTr | NA |  |
|  |  |  | Contrasts | Lo – Cm |  | **0.0473** |
|  |  |  |  | Lo – Co |  | 1.000 |
|  |  |  |  | Lo – Cr |  | 1.000 |
|  |  |  |  | Lo – Tr |  | NA |
|  |  |  |  | Cm – Co |  | 0.2189 |
|  |  |  |  | Cm – Cr |  | 1.000 |
|  |  |  |  | Cm – Tr |  | NA |
|  |  |  |  | Co – Cr |  | 1.000 |
|  |  |  |  | Co – Tr |  | NA |
|  |  |  |  | Cr - Tr |  | NA |
|  | Gills | BSD.log+  Lake+  BSD.log:Lake | Trends | BSD.log | **0.04454893 : 0.1757606** |  |
|  |  |  | Contrasts | Lo – Cm |  | **< 0.0001** |
|  |  |  |  | Lo – Co |  | **< 0.0001** |
|  |  |  |  | Lo – Cr |  | **< 0.0001** |
|  |  |  |  | Lo – Tr |  | 1.000 |
|  |  |  |  | Cm – Co |  | 1.000 |
|  |  |  |  | Cm – Cr |  | **< 0.0001** |
|  |  |  |  | Cm – Tr |  | **< 0.0001** |
|  |  |  |  | Co – Cr |  | **0.0002** |
|  |  |  |  | Co – Tr |  | **< 0.0001** |
|  |  |  |  | Cr - Tr |  | **< 0.0001** |
|  |  | CD.log+  Lake+  CD.log:Lake | Trends | CD.log | -0.1046101 : 0.02088965 |  |
|  |  |  | Contrasts | Lo – Cm |  | **< 0.0001** |
|  |  |  |  | Lo – Co |  | **< 0.0001** |
|  |  |  |  | Lo – Cr |  | **< 0.0001** |
|  |  |  |  | Lo – Tr |  | 1.000 |
|  |  |  |  | Cm – Co |  | 1.000 |
|  |  |  |  | Cm – Cr |  | **0.0002** |
|  |  |  |  | Cm – Tr |  | **0.0001** |
|  |  |  |  | Co – Cr |  | **0.0001** |
|  |  |  |  | Co – Tr |  | **< 0.0001** |
|  |  |  |  | Cr - Tr |  | **< 0.0001** |
|  |  | BS_Cest_D.log+  Lake+  BS_Cest_D.log:Lake | Trends | BS_Cest_D.log | **0.03151308 : 0.1648606** |  |
|  |  |  | Contrasts | Lo – Cm |  | **< 0.0001** |
|  |  |  |  | Lo – Co |  | **< 0.0001** |
|  |  |  |  | Lo – Cr |  | **< 0.0001** |
|  |  |  |  | Lo – Tr |  | 1.000 |
|  |  |  |  | Cm – Co |  | 1.000 |
|  |  |  |  | Cm – Cr |  | **< 0.0001** |
|  |  |  |  | Cm – Tr |  | **0.0001** |
|  |  |  |  | Co – Cr |  | **0.0002** |
|  |  |  |  | Co – Tr |  | **0.0001** |
|  |  |  |  | Cr - Tr |  | **< 0.0001** |
|  | Spleen | CD.log+  Lake+  CD.log:Lake | Trends | CD.log | **0.305347 : 0.5526565** |  |
|  |  |  | Contrasts | Lo – Cm |  | **< 0.0001** |
|  |  |  |  | Lo – Co |  | 1.000 |
|  |  |  |  | Lo – Cr |  | 1.000 |
|  |  |  |  | Lo – Tr |  | 1.000 |
|  |  |  |  | Cm – Co |  | **0.0033** |
|  |  |  |  | Cm – Cr |  | **< 0.0001** |
|  |  |  |  | Cm – Tr |  | **0.0020** |
|  |  |  |  | Co – Cr |  | 0.4219 |
|  |  |  |  | Co – Tr |  | 1.000 |
|  |  |  |  | Cr - Tr |  | 1.000 |
| LDH | Brain | CD.log+  Lake+  CD.log:Lake | Trends | CD.log:LakeLo | **0.000405 : 0.353** |  |
|  |  |  |  | CD.log:LakeCm | -0.176814 : 0.121 |  |
|  |  |  |  | CD.log:LakeCo | -0.459014 : 0.258 |  |
|  |  |  |  | CD.log:LakeCr | -1.572950 : 0.021 |  |
|  |  |  |  | CD.log:lakeTr | NA |  |
|  |  |  | Contrasts | Lo – Cm |  | 0.4873 |
|  |  |  |  | Lo – Co |  | 1.000 |
|  |  |  |  | Lo – Cr |  | 0.1360 |
|  |  |  |  | Lo – Tr |  | NA |
|  |  |  |  | Cm – Co |  | 1.000 |
|  |  |  |  | Cm – Cr |  | 0.4210 |
|  |  |  |  | Cm – Tr |  | NA |
|  |  |  |  | Co – Cr |  | 0.7696 |
|  |  |  |  | Co – Tr |  | NA |
|  |  |  |  | Cr - Tr |  | NA |
|  | Heart | BSD.log+  Lake+  BSD.log:Lake | Trends | BSD.log | **0.1964515 : 0.5004977** |  |
|  |  |  | Contrasts | Lo – Cm |  | **< 0.0001** |
|  |  |  |  | Lo – Co |  | **< 0.0001** |
|  |  |  |  | Lo – Cr |  | **< 0.0001** |
|  |  |  |  | Lo – Tr |  | 1.000 |
|  |  |  |  | Cm – Co |  | 0.0716 |
|  |  |  |  | Cm – Cr |  | 1.000 |
|  |  |  |  | Cm – Tr |  | **< 0.0001** |
|  |  |  |  | Co – Cr |  | 0.0575 |
|  |  |  |  | Co – Tr |  | **< 0.0001** |
|  |  |  |  | Cr - Tr |  | **< 0.0001** |
|  |  | CD.log+  Lake+  CD.log:Lake | Trends | CD.log | -0.1944144 : 0.07844882 |  |
|  |  |  | Contrasts | Lo – Cm |  | **< 0.0001** |
|  |  |  |  | Lo – Co |  | **< 0.0001** |
|  |  |  |  | Lo – Cr |  | **< 0.0001** |
|  |  |  |  | Lo – Tr |  | 1.000 |
|  |  |  |  | Cm – Co |  | 0.1484 |
|  |  |  |  | Cm – Cr |  | 1.000 |
|  |  |  |  | Cm – Tr |  | **< 0.0001** |
|  |  |  |  | Co – Cr |  | 0.1819 |
|  |  |  |  | Co – Tr |  | **< 0.0001** |
|  |  |  |  | Cr - Tr |  | **< 0.0001** |
|  | Gills | BSD.log+  Lake+  BSD.log:Lake | Trends | BSD.log | **0.1400155 : 0.4215580** |  |
|  |  |  | Contrasts | Lo – Cm |  | **< 0.0001** |
|  |  |  |  | Lo – Co |  | **< 0.0001** |
|  |  |  |  | Lo – Cr |  | **< 0.0001** |
|  |  |  |  | Lo – Tr |  | 0.4222 |
|  |  |  |  | Cm – Co |  | 1.000 |
|  |  |  |  | Cm – Cr |  | 1.000 |
|  |  |  |  | Cm – Tr |  | **< 0.0001** |
|  |  |  |  | Co – Cr |  | 1.000 |
|  |  |  |  | Co – Tr |  | **< 0.0001** |
|  |  |  |  | Cr - Tr |  | **< 0.0001** |
|  | Spleen | BSD.log+  Lake+  BSD.log:Lake | Trends | BSD.log | **0.2876113 : 0.5588732** |  |
|  |  |  | Contrasts | Lo – Cm |  | **< 0.0001** |
|  |  |  |  | Lo – Co |  | **< 0.0001** |
|  |  |  |  | Lo – Cr |  | **< 0.0001** |
|  |  |  |  | Lo – Tr |  | 0.2308 |
|  |  |  |  | Cm – Co |  | 0.4922 |
|  |  |  |  | Cm – Cr |  | **0.0010** |
|  |  |  |  | Cm – Tr |  | **< 0.0001** |
|  |  |  |  | Co – Cr |  | **< 0.0001** |
|  |  |  |  | Co – Tr |  | **< 0.0001** |
|  |  |  |  | Cr - Tr |  | **< 0.0001** |
|  |  | BS_Cest_D.log+  Lake+  BS_Cest_D.log:Lake | Trends | BS_Cest_D.log | **0.3132586 : 0.5847214** |  |
|  |  |  | Contrasts | Lo – Cm |  | **< 0.0001** |
|  |  |  |  | Lo – Co |  | **< 0.0001** |
|  |  |  |  | Lo – Cr |  | **< 0.0001** |
|  |  |  |  | Lo – Tr |  | 0.6568 |
|  |  |  |  | Cm – Co |  | 0.4258 |
|  |  |  |  | Cm – Cr |  | **0.0025** |
|  |  |  |  | Cm – Tr |  | **< 0.0001** |
|  |  |  |  | Co – Cr |  | **< 0.0001** |
|  |  |  |  | Co – Tr |  | **< 0.0001** |
|  |  |  |  | Cr - Tr |  | **< 0.0001** |
